# Ice Finder: Few-Shot Learning for Non-Vitrified Ice Segmentation

**DOI:** 10.1101/2024.08.05.606577

**Authors:** Alma Vivas-Lago, Daniel Castaño-Díez

**Affiliations:** Basque Centre for Biophysics (CSIC-UPV/EHU), Bilbao, Spain

## Abstract

This study introduces Ice Finder, a novel tool for quantifying crystalline ice in tomography, filling a crucial gap in existing methodologies. We establish the first application of the meta-learning paradigm to tomography, demonstrating that various tomographic tasks across datasets can be unified under a single meta-learning framework. Our approach utilizes few-shot learning to enhance domain generalization and adaptability to domain shifts, facilitating rapid adaptation to new datasets with minimal examples. Ice Finder’s performance is evaluated on a comprehensive set of in situ datasets from EMPIAR, proving its ease of use and fast processing capabilities, with inference times in the milliseconds. This tool not only accelerates workflows but also enhances the precision of structural studies in structural biology.

## Introduction

In the realm of structural biology, cryo-electron tomography (cryo-ET) has become indispensable for capturing detailed molecular landscapes of cells and tissues under conditions that closely mimic their natural state^1^. Historically significant, electron microscopy launched modern cell biology over sixty years ago by enabling the study of cellular ultrastructure. Today, cryo-ET extends this legacy by bridging the divide between molecular and cellular structural studies, providing subnanometer resolution within the cellular context^2^.

At the core of this technique is the process of vitrification, which preserves biological structures by rapidly freezing water into amorphous or vitreous ice^3^. This method avoids the formation of structured ice that could disrupt and distort cellular architecture, ensuring that samples retain a state nearly indistinguishable from their natural, hydrated condition^4^.

Traditionally, the stringent requirements of cryo-ET—such as precise control over sample thickness^5^ and the use of advanced preparation methods like cryo-sectioning^6,7^ or focused ion beam (FIB) milling^8–10^, limited its application to a few specialized labs^11–19^. These methods are crucial not only for achieving ultra-thin sections necessary for clear imaging but also for the meticulous optimization of cryoprotectants^20^, tailored specifically to each sample to prevent ice crystal formation^21,22^.

However, the field is experiencing a significant transformation. Advances in technology and procedural enhancements^23^, including the integration of light microscopes within FIB chambers^24–27^ and the automation of sample preparation^28,29^ and imaging^30–32^, have made cryo-ET more accessible. The availability of user-friendly, commercial equipment and the standardization of methods have broadened the technique’s appeal, supported by a growing network of specialized facilities that offer both essential tools and expert guidance, solidifying its role as a fundamental technique in structural biology^1^.

This democratization has expanded its use among novices and experts alike, facilitating complex workflows such as those integrating cryo-correlative light and electron microscopy (cryo-CLEM)^33^ to identify specific macromolecular complexes within crowded cellular environments^34^. Despite these advancements, challenges in sample preparation remain, particularly with vitrification. As the field pushes towards studying increasingly complex macromolecular assemblies^35^, the steps required to ensure complete vitrification frequently fall short^36^. This issue is compounded by prolonged workflows that increase the likelihood of crystal nucleation^37^. The subsequent growth of these crystals and further devitrification lead to significant phase changes, producing dramatic diffraction contrasts that can obscure and distort the structures of interest, ultimately complicating the interpretation of micrographs due to artifact-driven image formation^38^.

Despite the emergence of numerous new software tools aimed at enhancing image analysis and data quality^39^, a crucial gap remains: there is currently no tool that can effectively quantify the presence of structured ice within tomographic samples. While detection is straightforward in single-particle cryo-EM, where tools like CryoSPARC^40^ efficiently remove problematic micrographs, cryo-ET encounters greater challenges. In single-particle cryo-EM^41^, each sample is imaged only once, allowing the full electron dose to be utilized. This maximizes the signal-to-noise ratio, generating high-resolution datasets containing hundreds of thousands to millions of particle images of the biological molecule of interest, providing an ideal “portrait” of the target molecule^42–45^. In contrast, tomography^46^ acquires a series of images at different tilts, fractionating the dose across multiple images. This method often involves cellular or tissue environments where the surroundings are more crowded and complex^47^. Consequently, cryo-ET faces lower signal-to-noise ratios and sometimes inadequate pixel sizes, complicating the detection of diffraction peaks (see Fig. 1). Addressing this gap with the introduction of an automated tool capable of quantifying structured ice presence in in situ workflows is critical. Such a tool would diminish the need for manual inspection of hundreds of tomograms and significantly accelerate structural studies, better meeting the demands of modern biological research^48^.

**Figure 1.**
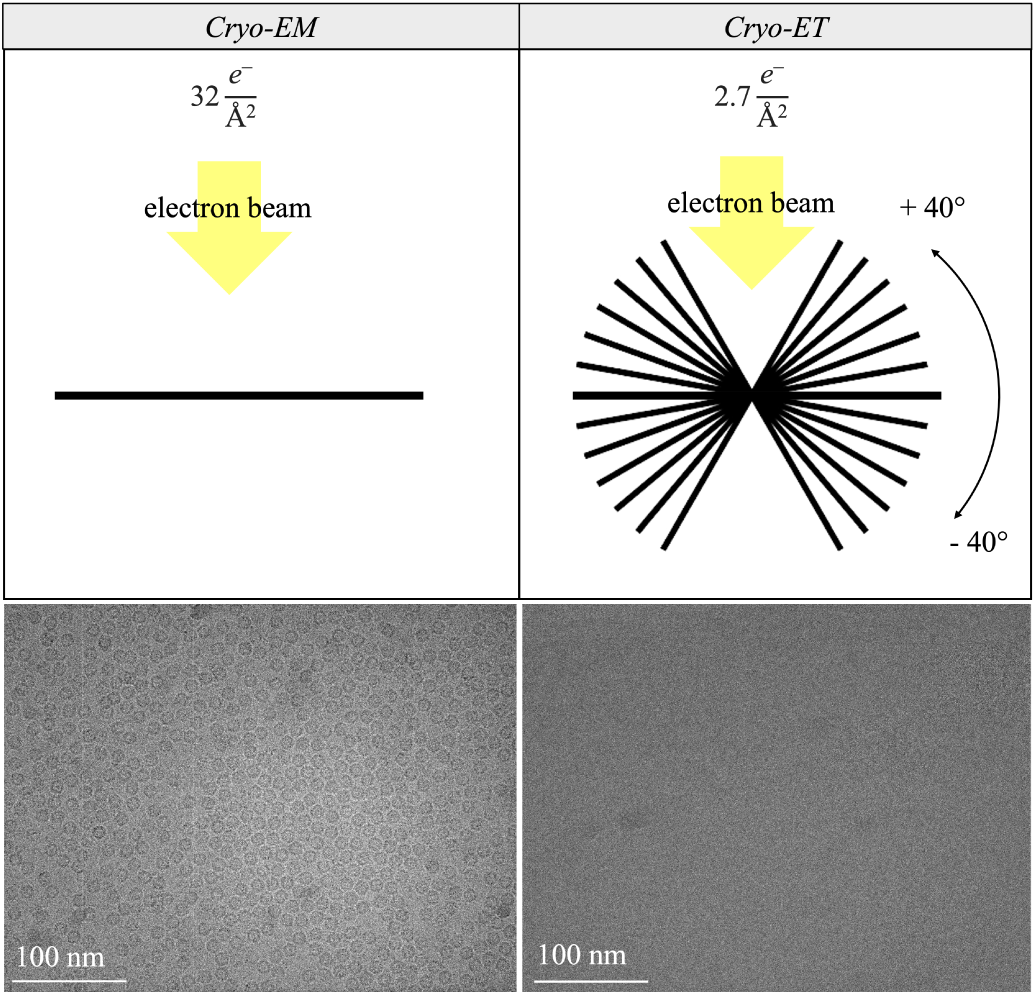
Comparison of data collection strategies: Single-particle cryo-EM (left) with a single high-dose image (32 e*^−^* Å*^−^*^2^), and cryo-ET (right) with multiple lower-dose tilt images (2.7 e*^−^* Å*^−^*^2^). The sample is Apoferritin from the EMPIAR-10491 frame and tilt-series data collected from the same grid square under identical conditions.

The inherent complexity and variability in biological samples pose significant challenges for standardizing image analysis in tomography. Each sample exhibits unique characteristics, leading to substantial intra- and inter-sample variability. This reality underscores the need for systems that can quickly adapt to new datasets and learn effectively from just a few examples.

Encouraging the ability of a system to learn from a limited number of examples is referred to in the literature as *few-shot learning* (FSL). This property can be achieved by enhancing with prior knowledge three building blocks of the system: data, model, and algorithm^49^. The overall objective of the modifications is to make the empirical risk minimizer reliable again.

Typically, when data is enhanced, the goal is to increase the number of instances in the training dataset. This can be achieved through various methods: learning a transformation function to apply to the original training samples, developing a predictor based on the original training data which is then used on a weakly labeled or unlabeled dataset, or creating an aggregator function that modifies samples from similar datasets.

Conversely, solutions that focus on modifying the model aim to use prior knowledge to constrain the hypothesis space. Several strategies have been developed for this purpose, known in the machine learning literature as multi-task learning, embedding learning, learning with external memory, and generative modeling. Delving into each of them is out of the scope of this manuscript, the interested reader is referred to^49^.

Finally, prior knowledge is utilized to modify the search strategy within the hypothesis space to identify the parameters of the best hypothesis. Approaches like these impact the parameter discovery process by either providing well-initialized parameters or teaching the optimizer to navigate the search steps effectively. A well-known technique that falls within this category is *transfer learning*, where an initial parameter set is learned from a different task and then refined using training data specific to the current task. Since this technique often does not work well when the target dataset is small, our study explores an alternative approach known as Model-Agnostic Meta-Learning (MAML), which refines *meta-learned* parameters for effective few-shot learning applications^50^.

Meta-learning, often described as “learning-to-learn”^51^, leverages the experience accumulated over multiple learning episodes, each encompassing a distribution of related tasks. This methodology aims to enhance the efficiency and effectiveness of future learning processes. By distilling^52,53^ insights from these episodes, meta-learning improves both data and computational efficiency, aligning more closely with human and animal learning, where learning strategies evolve over both individual lifetimes and evolutionary timescales^54–57^.

Historically, neural network meta-learning has a rich lineage, but its potential has only recently catalyzed a surge in research within the deep learning community^58–62^. This resurgence is driven by meta-learning’s promise to address key limitations of traditional deep learning, particularly through enhanced data efficiency and superior knowledge transfer capabilities^57^.

Meta-learning excels in multi-task environments, where task-agnostic knowledge derived from a diverse array of tasks can be harnessed to expedite learning in new, related tasks^63^. This approach aligns with the perspective of using meta-learning to mitigate the ‘no free lunch’ theorem^64^ by optimizing the inductive biases that best suit specific problem domains. Beyond the traditional scope, contemporary neural network meta-learning is formulated as end-to-end learning of explicitly defined objective functions, thereby refining the learning processes through these tailored goals^57^.

While transfer learning^65^ is traditionally favored for its ability to utilize prior knowledge from related tasks to enhance performance on new tasks, our study also considers the broader spectrum of learning strategies, including meta-learning. Both approaches will be explored to determine which is more effective at meeting our objectives in the context of limited data availability.

We present a general schematic view of our learning framework tailored to few-shot learning. This illustration, depicted in Fig. 2, demonstrates the process of fine-tuning initially learned parameters based on a few labeled examples from the target domain. Our aim is to identify the most efficient method—be it transfer learning or meta-learning—for achieving sample-efficient learning, domain generalization, and robustness to domain shifts within the challenging environment of tomography.

**Figure 2.**
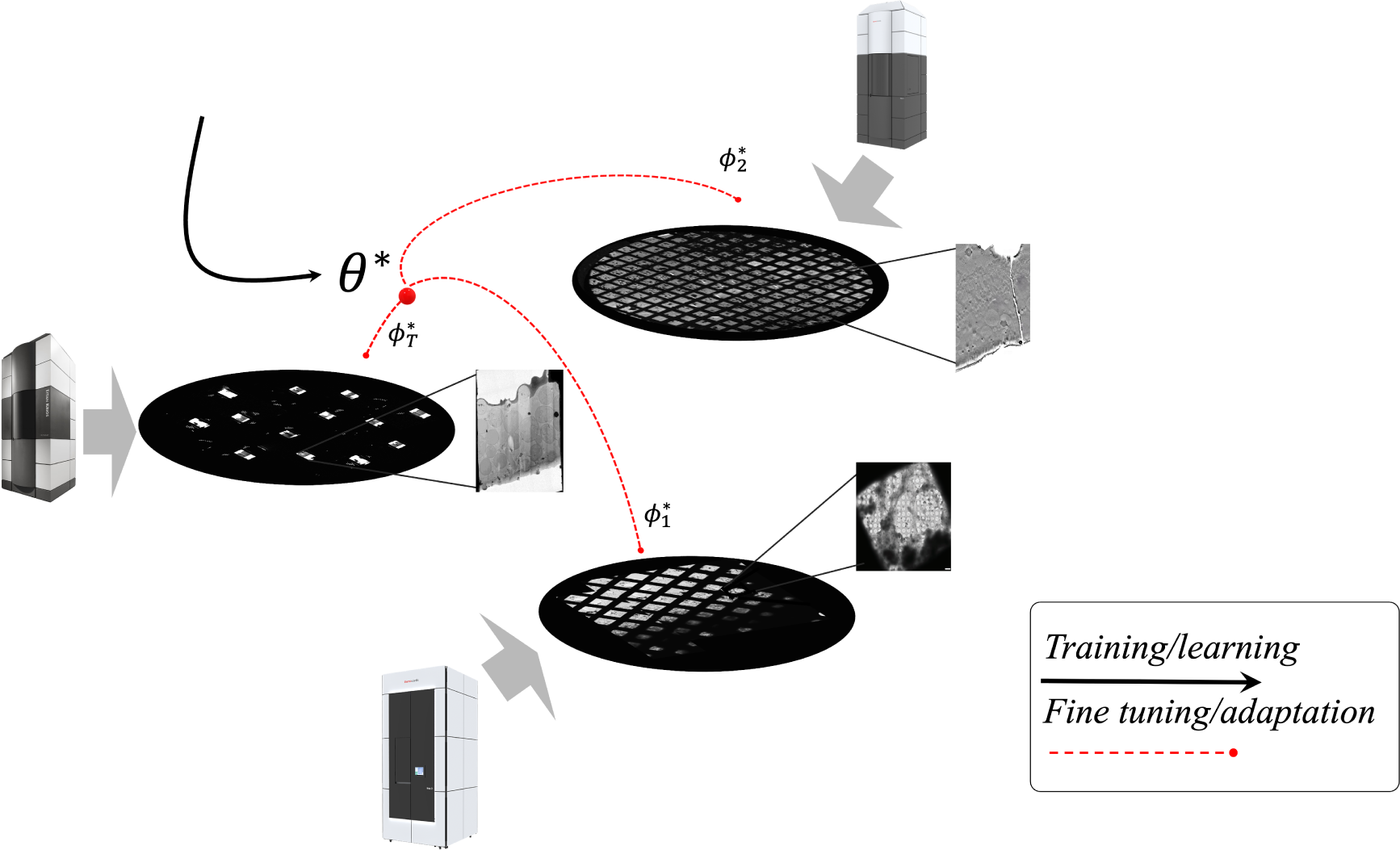
Schematic view of the few-shot learning framework applied to tomographic analysis. This diagram illustrates the process of adapting initial optimized weights *θ^∗^* to new datasets with limited examples, resulting in optimized task-specific parameters 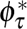. This visualization underscores the complexity of tomographic datasets, which can encompass thousands of micrographs across multiple grids, each with unique sample contents and configurations. The diversity and scale of these datasets pose significant computational challenges, necessitating robust few-shot learning strategies for effective adaptation and learning.

Our main contributions are threefold: (1) we developed a new tool to precisely quantify crystalline ice, addressing a significant gap in current methodologies; (2) we demonstrated the first proof of concept of applying the meta-learning paradigm to tomography, showing that tomographic tasks across datasets can be unified under a single meta-learning framework; and (3) we evaluated the efficacy of our Ice Finder tool for few-shot segmentation using a comprehensive set of publicly available in situ datasets.

## Proposed method

### Segmentation model

In this study, we employ the Feature Pyramid Network (FPN) architecture for the segmentation task. The choice of FPN over other architectures such as U-Net is driven by its superior handling of objects at different scales. The general architectures of both U-Net and FPN are illustrated in Fig. 3.

**Figure 3.**
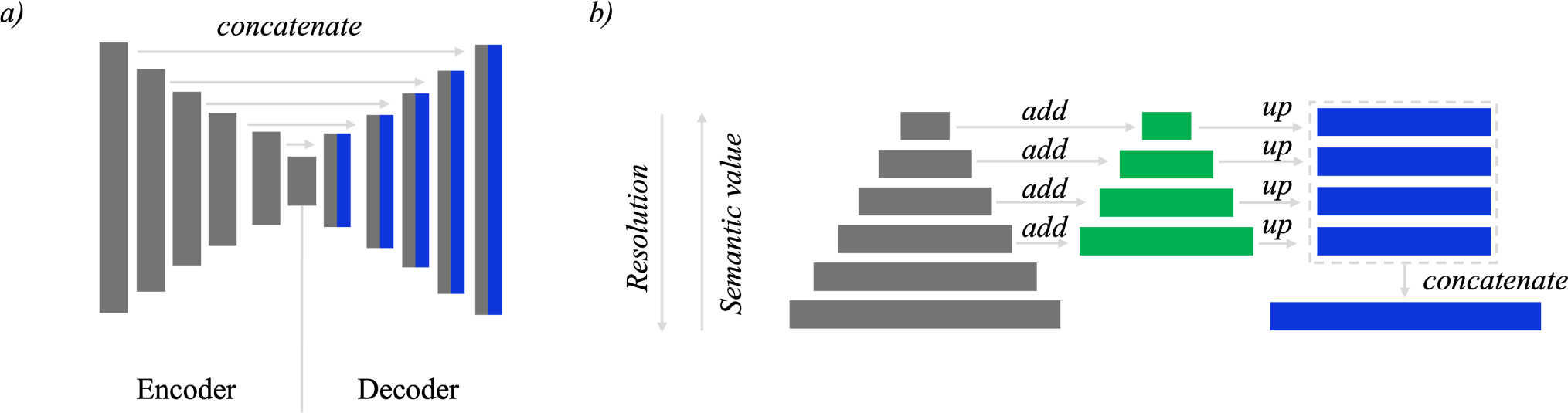
The general architecture of the (a) U-Net and (b) FPN model. Adapted from^66^.

The U-Net architecture, widely recognized for its success in semantic segmentation tasks^67^, follows a symmetric encoder-decoder structure. This design includes multiple upsampling layers and skip connections that concatenate features from the encoder to the decoder, enhancing the ability to capture fine details. However, in our scenario, the regions of interest within each micrograph vary significantly in size, which necessitates a model that can efficiently manage such scale variations.

FPN^68^ leverages the inherent multi-scale, pyramidal hierarchy of deep convolutional networks to construct feature pyramids with minimal additional computational cost. This architecture consists of a top-down pathway combined with lateral connections, which enables the creation of high-level semantic feature maps at multiple scales. The result is a robust representation that maintains strong semantic features across different resolution levels, making it particularly effective for detecting and segmenting objects of varying sizes within the same image.

## Backbone Architectures

In the context of fine-tuning, the choice of backbone architecture is crucial^69^ due to its impact on gradient behavior and training stability. The ResNet family is renowned for its efficacy in fine-tuning tasks, often leading to well-behaved gradients^70^. Despite their robustness, ResNet architectures can be challenging to train. To address this, we explore DenseNet, a natural extension of ResNet, which offers enhanced feature reuse and efficiency^71^.

### Residual Network

The key idea of Residual Networks is the use of identity mappings. Formally, a building block is to be defined as:

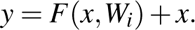

where *x* and *y* are the input and output vectors of the layers considered. The function *F*(*x,W_i_*) represents the residual mapping to be learned^72^.

In the revisited version, the propagation formulations allow signals (information) to be directly propagated from one block to any other block if an additive identity transformation is used. Coupling this idea with the original model’s skip connections and post-addition activation yields promising results^73^.

### DenseNet

DenseNet builds on ResNet, offering a natural extension with unique properties. Instead of adding the input to the mapping function, it concatenates it. This approach allows each layer to have direct access to the feature maps of all its preceding layers, creating a “collective knowledge” of features within its blocks. This architecture facilitates feature reuse, resulting in higher parameter efficiency. Furthermore, DenseNet networks are considered easier to train^74^ due to the improved gradient flow through the network.

## Few-shot Learning Framework

### Transfer Learning

Transfer learning tasks consist of a source and a target dataset, differing in terms of their underlying distribution.

Stated formally^75^:

*Given a source domain D_S_ with input data X_S_, a corresponding source task T_S_ with labels Y_S_, as well as a target domain D_T_ with input data X_T_ and a target task T_T_ with labels Y_T_, the objective of transfer learning is to learn the target conditional probability distribution P_T_* (*Y_T_ |X_T_*) *with the information gained from D_S_ and T_S_ where D_S_ ≠ D_T_ or TS ≠ T_T_*.

In this regard, in a fully supervised setting, there are two possible scenarios: domain adaptation, task containing a small amount of additional labeled target data, or domain generalization task, where the access is restricted to labeled source data.

### Meta-learning

Meta-learning strategies differ significantly in how they utilize meta-knowledge, which determines the aspects of a learning strategy that are adaptive versus those that are fixed^57^. In our research, we employ optimization-based meta-learning methods, valued for their robust consistency and superior generalization capabilities over black-box or non-parametrized approaches^50^. Notably, Model-Agnostic Meta-Learning (MAML) stands out for maintaining inductive bias, crucial for expressive model performance without a loss in generality^76^.

MAML, an exemplar of optimization-based meta-learning, employs a computation graph embedded with gradient operations, making it distinct from conventional single-level optimization frameworks. This method requires a meta-dataset—a collection of diverse yet related datasets^63^, supporting bi-level optimization.

**Key Concept:** Within MAML, tasks are treated as independent data points drawn from a shared task distribution *p*(*T*), assumed to be independently and identically distributed (i.i.d.).

The meta-learning framework is formalized through key components:

**Task** *τ*: Defined by its unique support and query sets.

**Support Set** 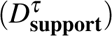: Data subset for model parameter adaptation.

**Query Set** 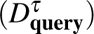: Data subset for evaluating model adaptation performance.

**Task Distribution** *p*(*ℐ*): The probabilistic distribution from which tasks are sampled.

**Model Parameters** *θ* : Initial shared parameters optimized across tasks during meta-training.

**Task-Specific Parameters** *φ_τ_* : Adapted parameters specific to each task.

To contrast with traditional model training, where parameters *θ* are optimized through gradient descent to perform well on a single dataset (Algorithm 1), MAML optimizes initial model parameters *θ* to facilitate rapid adaptation across a spectrum of tasks (Algorithm 2). This dual-level optimization process, especially the adaptation in the meta-learning scenario, involves second-order gradient computations. Specifically, the gradient of the loss with respect to *θ* is contingent upon the gradients of the task-specific adapted parameters *φ_τ_*, making the method computationally expensive but highly effective for generalization across varied tasks.

By optimizing the learning process itself, MAML ensures robust generalization across different, but related tasks, thereby broadening the scope and applicability of the learned models in real-world scenarios.

#### Algorithm 1 Regular Gradient Descent

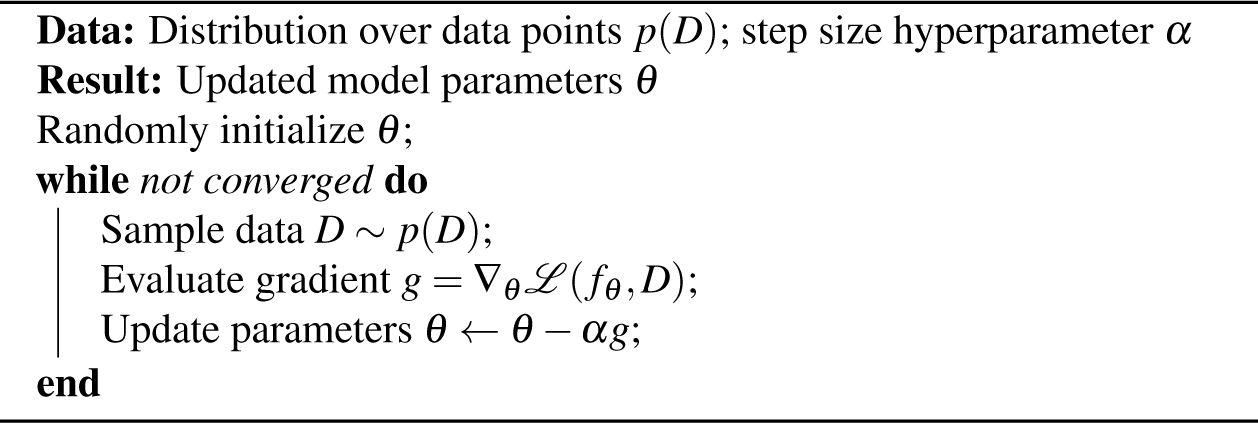

#### Algorithm 2 Model-Agnostic Meta-Learning (MAML)

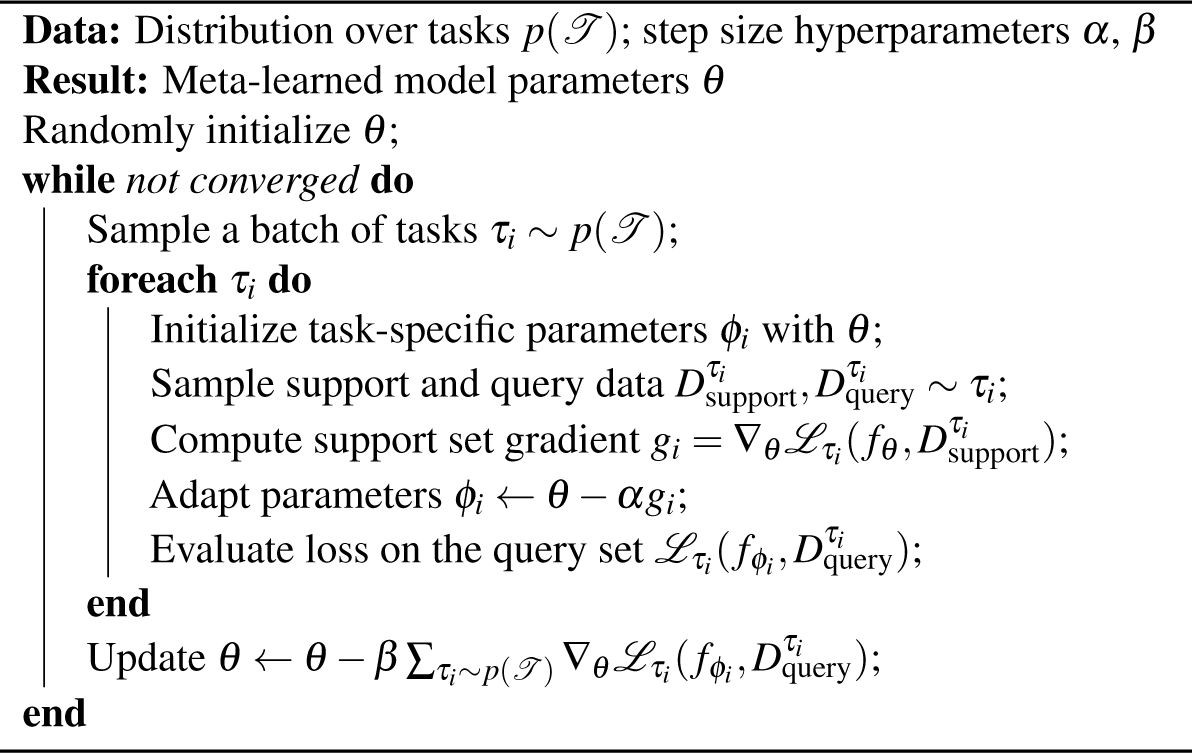

#### Task Design

In meta-learning, tasks must share a common structure, akin to the principle in transfer learning^77^. This shared structure can be either abstract or concrete. In our context, the structure refers to the optical characteristics of ice in its various phases, as shown in Fig. 4. We primarily deal with hexagonal ice (I*_h_*), cubic ice (I*_c_*), and, in the majority of cases, stacking-disordered ice I (ice (I_sd_))^78^. Studies have shown that distinguishing between cubic ice and stacking-disordered ice, which is a mixture of cubic and hexagonal sequences, can be challenging^79^. Our preliminary experiments, indicated by diffuse diffraction patterns, confirm the presence of both phases (results not shown). Typically, poor vitrification from insufficient rapid freezing leads to hexagonal ice formation, while cubic ice formation suggests that an initially vitreous sample has warmed and subsequently devitrified^80^. These findings indicate that in many cases, poor initial vitrification followed by warming leads to the formation of stacking-disordered ice.

**Figure 4.**
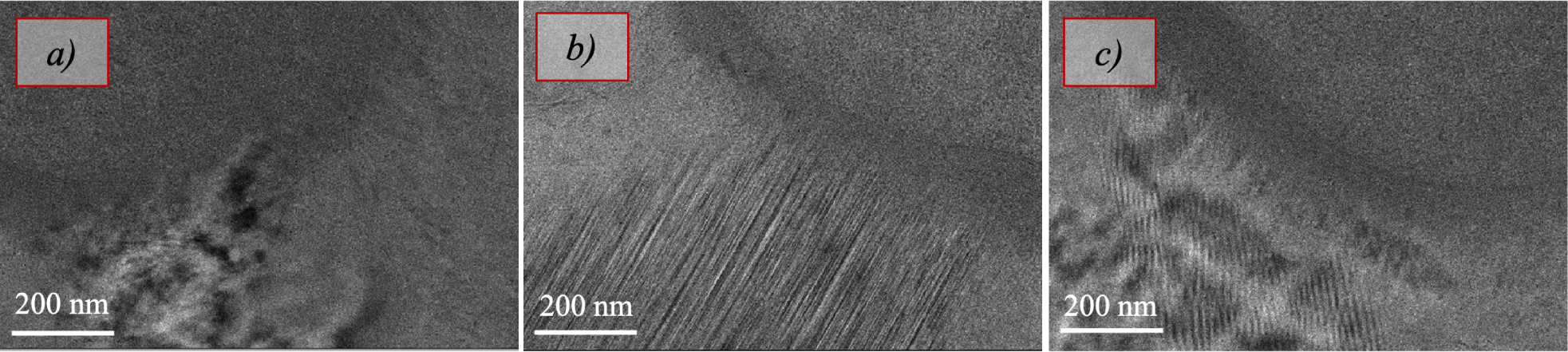
Crystalline phases of ice: hexagonal (I*_h_*) (a), cubic (I*_c_*) (b), and stacking-disordered ice (I_sd_) (c).

To design a task, we conceptualize it as a collection of micrographs—projections or images recorded from the microscope—from the same dataset, ensuring that they share the same microscope parameters/configuration, sample content, and preparation. For simplicity, we denote the dataset as a single grid, although it may encompass multiple grids. Each micrograph in a task comes from a different lamella—a thin slice of the sample—different tilt series, and different projection. This design ensures that the model learns the common features of crystalline ice while being exposed to diverse views and conditions, thus mimicking the behavior we aim to reproduce at meta-test time (see Fig. 5).

**Figure 5.**
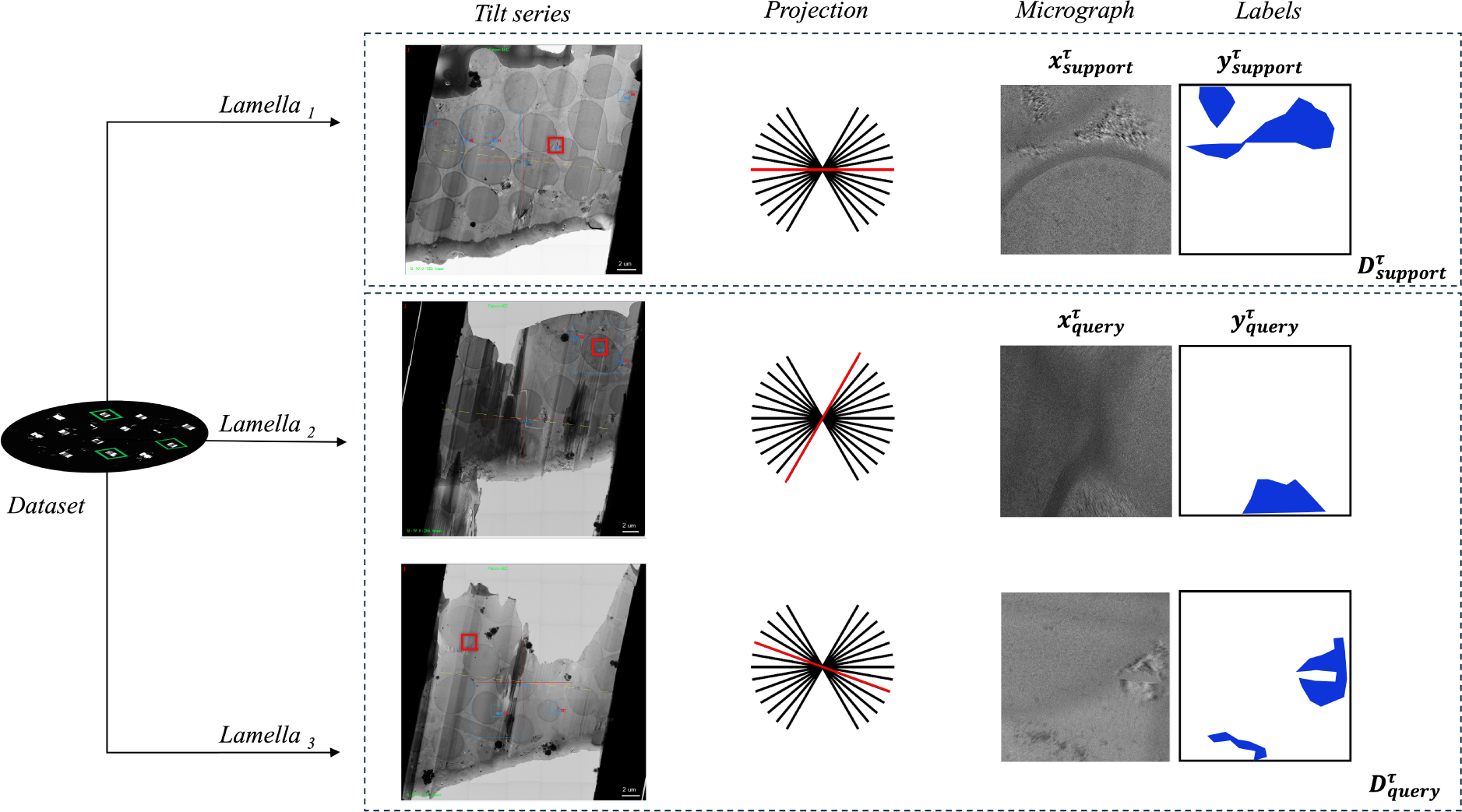
Illustration of task design for MAML. Each task is derived from a dataset consisting of micrographs captured under consistent microscope parameters and sample preparation conditions. The tasks include images from different lamellae, tilt series, and projections, divided into support sets 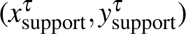 for model adaptation and query sets 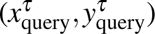 for inner loop performance evaluation. While the goal remains semantic segmentation, variations in data distribution make each task distinct. This division allows the model to learn generalizable features across different but related tasks.

In a supervised learning setting, each task is split into a support set and a query set. The support set 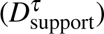 is used to adapt the model parameters to the specific task *τ*, while the query set 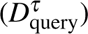 is used to evaluate the performance of the adapted model. This approach ensures that the model is trained to generalize well across different but related tasks, leveraging the shared structure of the data.

## Results

This study leverages datasets from EMPIAR (Electron Microscopy Public Image Archive)^81^, hosted by EMBL-EBI. This public repository offers a large variety of datasets from different biological samples collected by EM, being customarily used for development and validation in this area. Our dataset selection, detailed in Table 1, was strategically chosen to maximize diversity in biological characteristics, ensuring a varied test set as depicted in Fig. 6.

**Figure 6.**
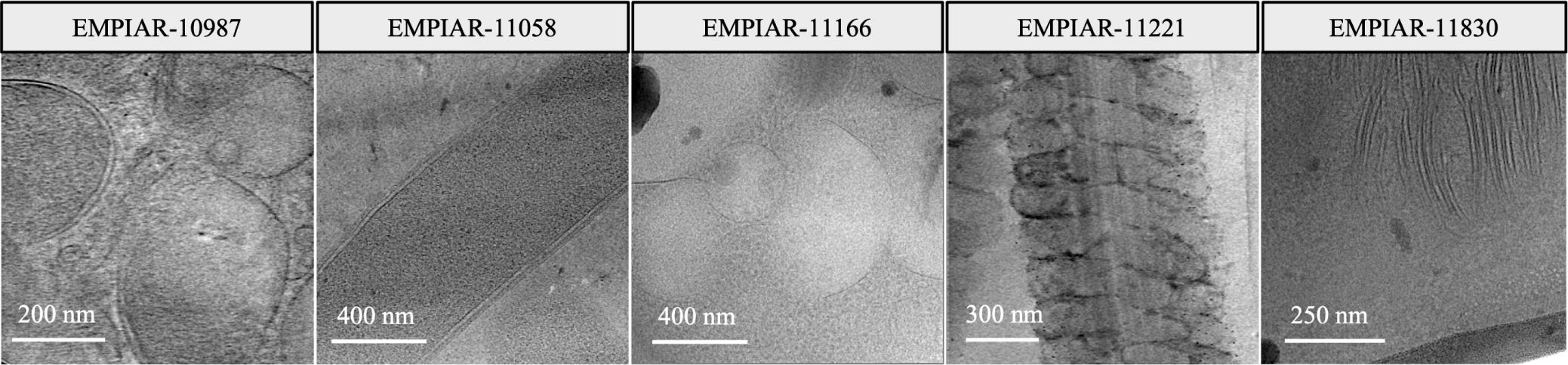
Overview of data used in the test set, showcasing the diversity of biological samples.

**Table 1.** Details of EMPIAR Entries used in the study.

| Entry ID | Pixel Size [Å] | Image Size [pixels] | Sample |
| --- | --- | --- | --- |
| EMPIAR-10987 | (1.64, 1.64) | (4092, 5760) | 80S ribosomes |
| EMPIAR-11058 | (3.52, 3.52) | (3712, 3712) | Thermoanaerobacter kivui |
| EMPIAR-11166 | (3.52, 3.52) | (3712, 3712) | Saccharomyces cerevisiae autophagy |
| EMPIAR-11221 | (3.46, 3.46) | (3710, 3838) | Mouse sperm flagella |
| EMPIAR-11830 | (1.90, 1.90) | (4096, 4096) | Chlamydomonas reinhardtii |

**Table 2.**
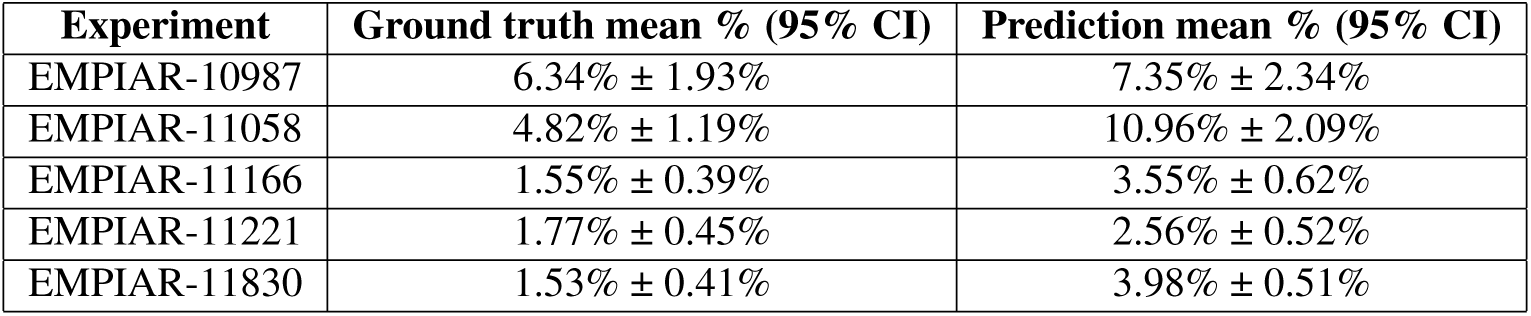
Quantitative analysis of domain generalization across different datasets, measured as percentages of non-vitrified areas correctly identified in inference mode.

| Experiment | Ground truth mean % (95% CI) | Prediction mean % (95% CI) |
| --- | --- | --- |
| EMPIAR-10987 | 6.34% ± 1.93% | 7.35% ± 2.34% |
| EMPIAR-11058 | 4.82% ± 1.19% | 10.96% ± 2.09% |
| EMPIAR-11166 | 1.55% ± 0.39% | 3.55% ± 0.62% |
| EMPIAR-11221 | 1.77% ± 0.45% | 2.56% ± 0.52% |
| EMPIAR-11830 | 1.53% ± 0.41% | 3.98% ± 0.51% |

For comprehensive context, a complete list of all in situ study samples considered is provided in the supplementary materials. This list includes samples not directly analyzed in our study but available for future research. We excluded datasets consisting only of tomograms, single tilt series, or those derived from alternative modalities such as X-ray tomography to maintain consistency and relevance to the objectives of our study.

From each dataset, tilt series exhibiting vitrification issues were identified. From the available items, three tilt series per dataset were selected. Out of these three processed tilt series, one was used for training in what is referred to as k-shot testing. The remaining two served as replicates to simulate a real-world scenario where the selected, fine-tuned, models were applied in inference mode to assess their performance. For a detailed overview of the implementation details, please see the supplementary material.

The workflow for utilizing the Ice Finder tool is designed to operate seamlessly within a Jupyter notebook^82^, where each step is compartmentalized into distinct notebook cells (see Fig. 7). Users begin by preprocessing their selected tilt series or micrographs (1 to 10 micrographs), which involves uploading the images through a simple interface that only requires specifying the path to the data and executing the preprocessing notebook cell. After preprocessing, users proceed to annotate the images using CVAT^83^, marking the crystalline regions with polygons. These annotations are then exported, and their paths are inputted back into the system for fine-tuning the model—this process is streamlined as hyperparameters are preset.

**Figure 7.**
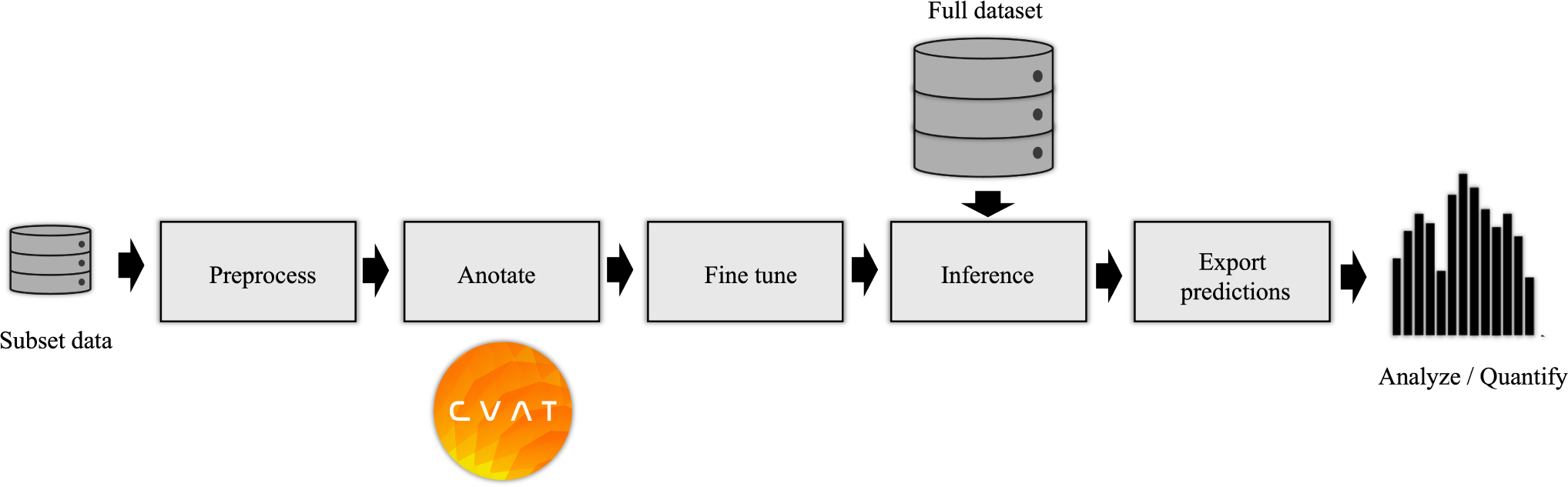
Workflow for using the Ice Finder tool: Users preprocess their data, annotate ice regions using CVAT, fine-tune the model with annotated examples from a pre-trained checkpoint on a large dataset, run inference on the full dataset, and export the analysis results. This streamlined process is integrated onto a Jupyter notebook, where each step is compartmentalized into distinct notebook cells.

A pre-trained checkpoint from training on a large dataset is provided. Based on the new sample, fine-tuning (k-shot) is performed, and post fine-tuning, the best model is automatically selected. Users run inference by providing the path to the complete dataset, initiating the model to analyze the full set of images. The final step in the workflow is the analysis phase, where users can either export the segmented masks or generate statistics about the percentage of ice present, aiding in the rapid assessment of vitrification quality across samples.

The following table presents a quantitative analysis of our models’ domain generalization capabilities by comparing the predicted percentages of non-vitrified areas against the ground truth across various datasets.

Although the Intersection over Union (IoU) metric is a standard evaluation metric in many segmentation tasks, we chose to use the percentage of non-vitrified areas for the final quantitative analysis due to its simplicity and higher relevance for end-users. This metric resonates well with the primary objective of our study, which is to quantify the extent of vitrification. By focusing on percentage measurements, we provide a straightforward and easily interpretable metric that aligns closely with the practical needs of the users in assessing vitrification quality.

This quantitative assessment provides a broad overview of model performance and sets the foundation for a detailed review of how each specific experiment adapted to its domain. The following deeper inspection aims to understand the adaptability and effectiveness of our approaches under diverse and challenging conditions typical in biological datasets. To this end, the performance of our baseline (vanilla transfer learning), MAML, and a random model as control, will be systematically compared.

### EMPIAR-10987

The models displayed consistent performance (see Fig. 8). This is attributed to effectively identifying the most prominent (i.e. salient), high-intensity ice patterns, while subtly revealing the lower frequency footprints as well. Figure 9 highlights their capability to detect pronounced patterns atop ethane blobs, although they occasionally underestimated the dimensions (see Fig. 9 first column). Notably, in the upper right of the extension, the ice was not detected, underscoring a limitation in recognizing less distinct features mixed with cubic phases (see Fig. 9, second column).

**Figure 8.**
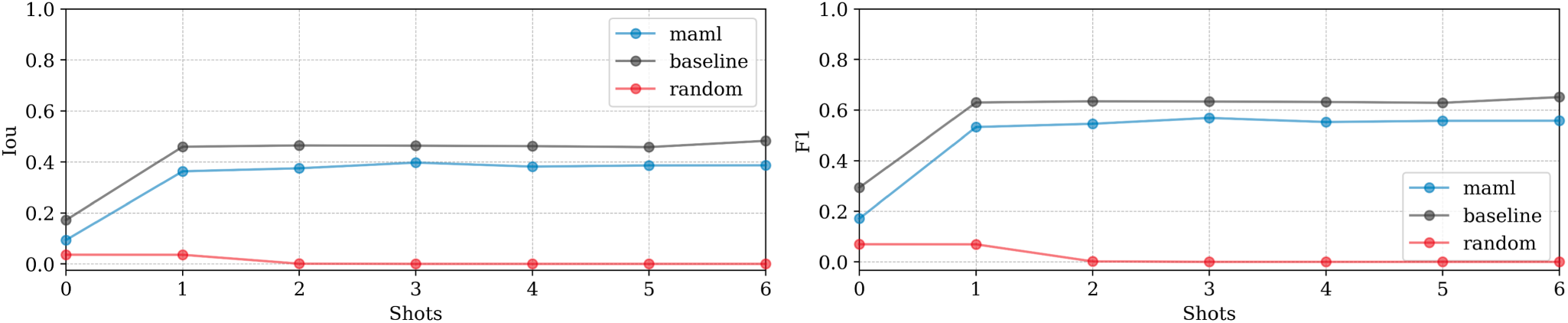
Domain adaptation performance: EMPIAR-10987. The graphs display the performance metrics of different models (MAML, baseline, and random) evaluated using Intersection over Union (IoU) and F1 scores across multiple k-shot scenarios. The consistency in performance reflects the models’ capability to generalize across various domains by effectively learning and adapting to new data distributions.

**Figure 9.**
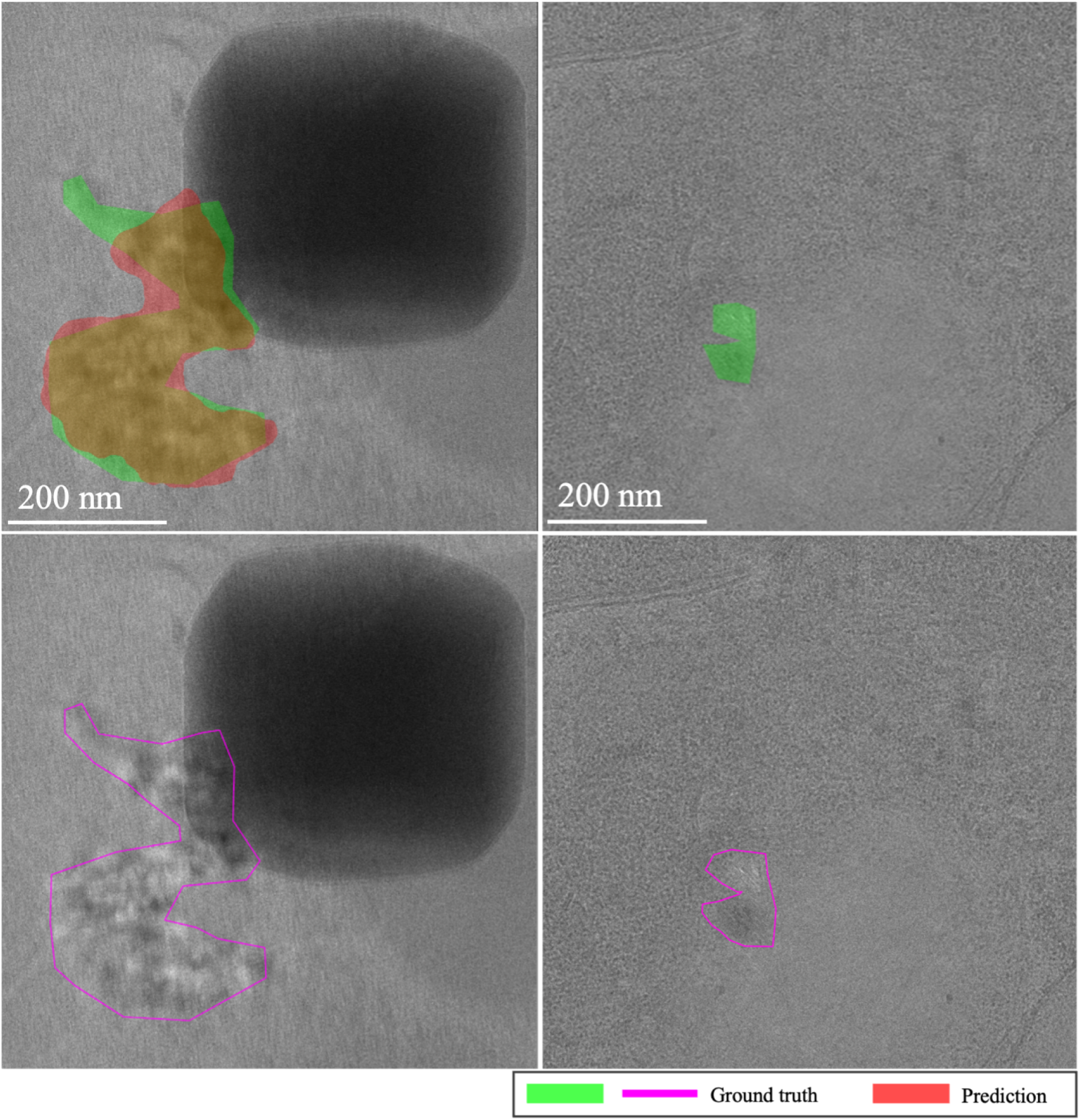
Main sources of error for EMPIAR-10987. The images illustrate the primary challenges faced by the models, such as underestimating the dimensions of ice regions (first column) and failing to detect less distinct ice features mixed with cubic phases (second column). Despite these challenges, the models successfully identified prominent ice patterns, demonstrating their robustness and adaptability in complex scenarios.

### EMPIAR-11058

Both models exhibited slight fluctuations in performance, synchronizing until the 4th k-shot where both achieved peak performance. Subsequently, at the 5th shot, the baseline model’s performance declined while MAML’s improved (see Fig. 10). This turning point corresponds to an image of a very thick lamella where the ice, appearing outside the bacterial structure and resembling a fluid-like, low-frequency pattern, offered a minor success for MAML—albeit still systematically below the baseline—highlighting its purported flexibility and robustness. The soft, fluid-like appearance, often shallow and weak, was typically unlabelled leading to false positive predictions as exemplified in Fig. 11. Another source of unexpected failures included misidentifications of bubbles caused by ion beam sample damage as depicted in Fig. 12.

**Figure 10.**
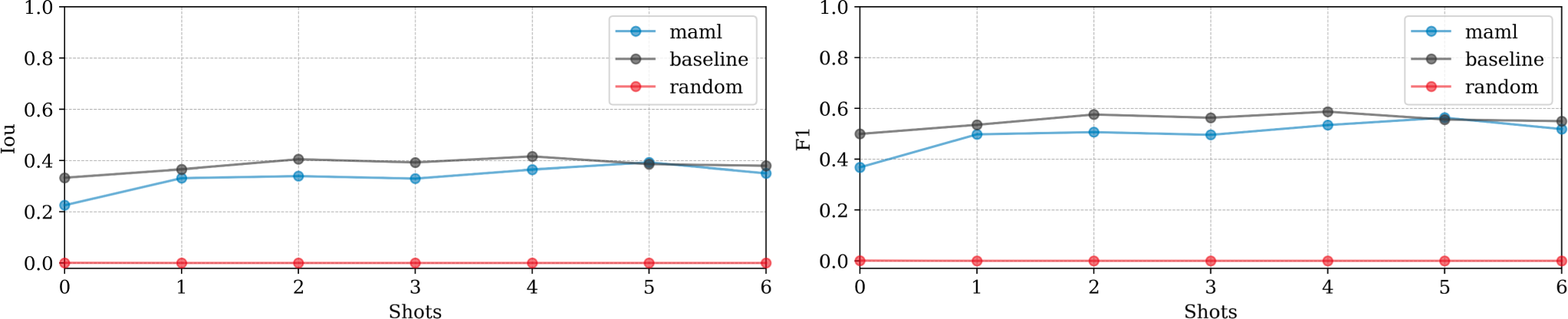
Domain adaptation performance: EMPIAR-11058. The graphs display the performance metrics of different models (MAML, baseline, and random) evaluated using Intersection over Union (IoU) and F1 scores across multiple k-shot scenarios. The performance variability reflects the models’ capability to generalize and adapt to new data distributions, with notable fluctuations corresponding to specific challenging samples.

**Figure 11.**
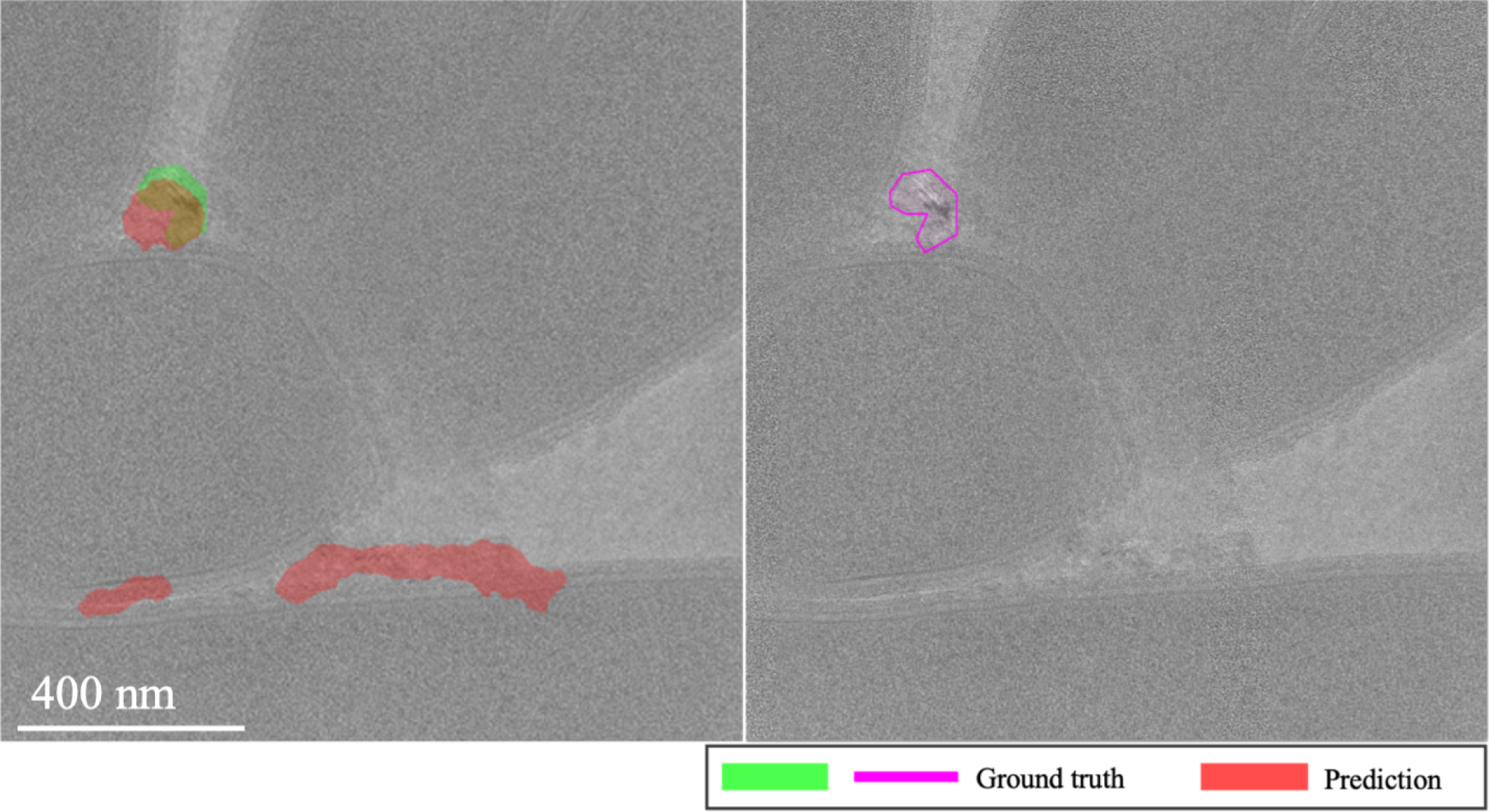
Sharp features misidentification in EMPIAR-11058. The images highlight the misidentification of soft, fluid-like ice features, often leading to false positive predictions due to the unlabelled shallow and weak appearance of these patterns.

**Figure 12.**
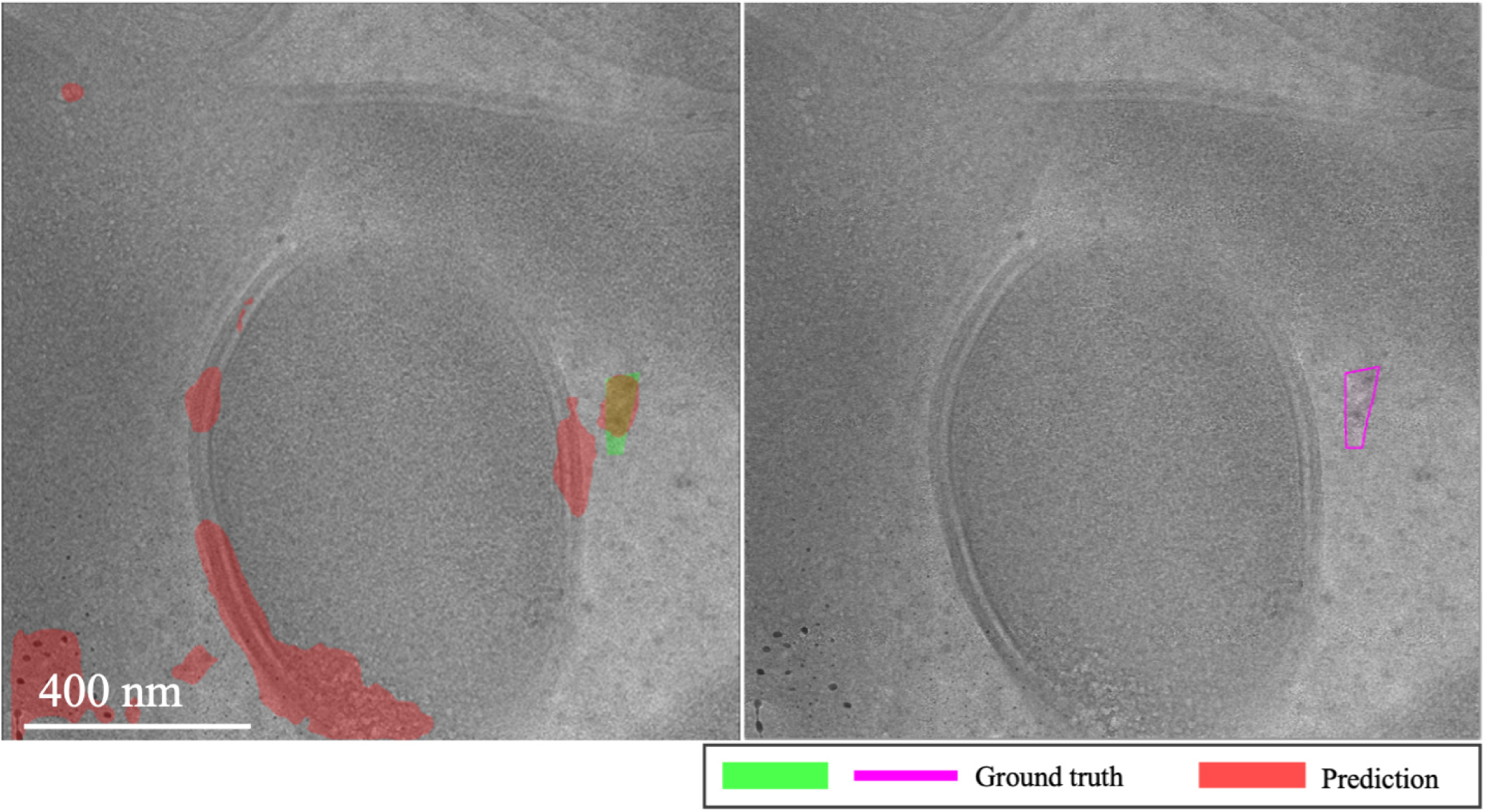
Ion beam damage misinterpretation in EMPIAR-11058. The images illustrate the misidentification of bubbles caused by ion beam sample damage, demonstrating a challenge in distinguishing these artifacts from actual ice features.

### EMPIAR-11166

Initial shots exhibited turbulence, with the baseline struggling to integrate labels and MAML adjusting to optimize task performance. By the third and fourth k-shots, performances of both models aligned closely, matching exactly at the fourth shot. Thereafter, the baseline model showed slight improvement with a minimal slope, while MAML’s performance diverged (see Fig. 13). This variability largely stemmed from inconsistencies between the models’ recognition of ice and the provided labels, which only marked prominently salient ice relative to the feature signal of the image. Consequently, predictions captured weaker yet existent areas affected by vitrification. This issue was apparent in sheet-like patterns (Fig. 14, first column) and isolated spots to small regions of low-frequency crystalline ice reflections (Fig. 14, second column).

**Figure 13.**
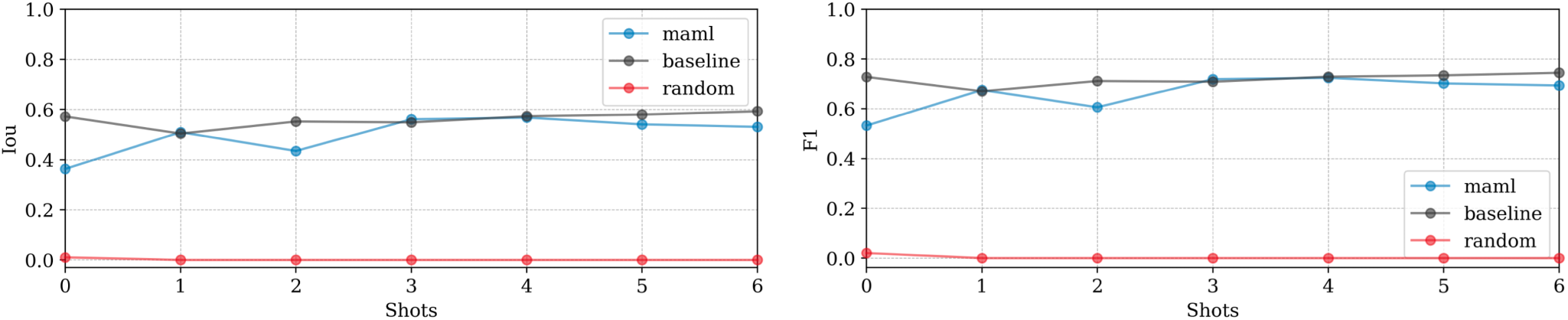
Domain adaptation performance: EMPIAR-11166. The graphs display the performance metrics of different models (MAML, baseline, and random) evaluated using Intersection over Union (IoU) and F1 scores across multiple k-shot scenarios. The performance shows minor variability due to label inconsistencies, despite the nearly constant performance reflecting the target’s similar distribution. This highlights the models’ capability to adjust and generalize across various domains.

**Figure 14.**
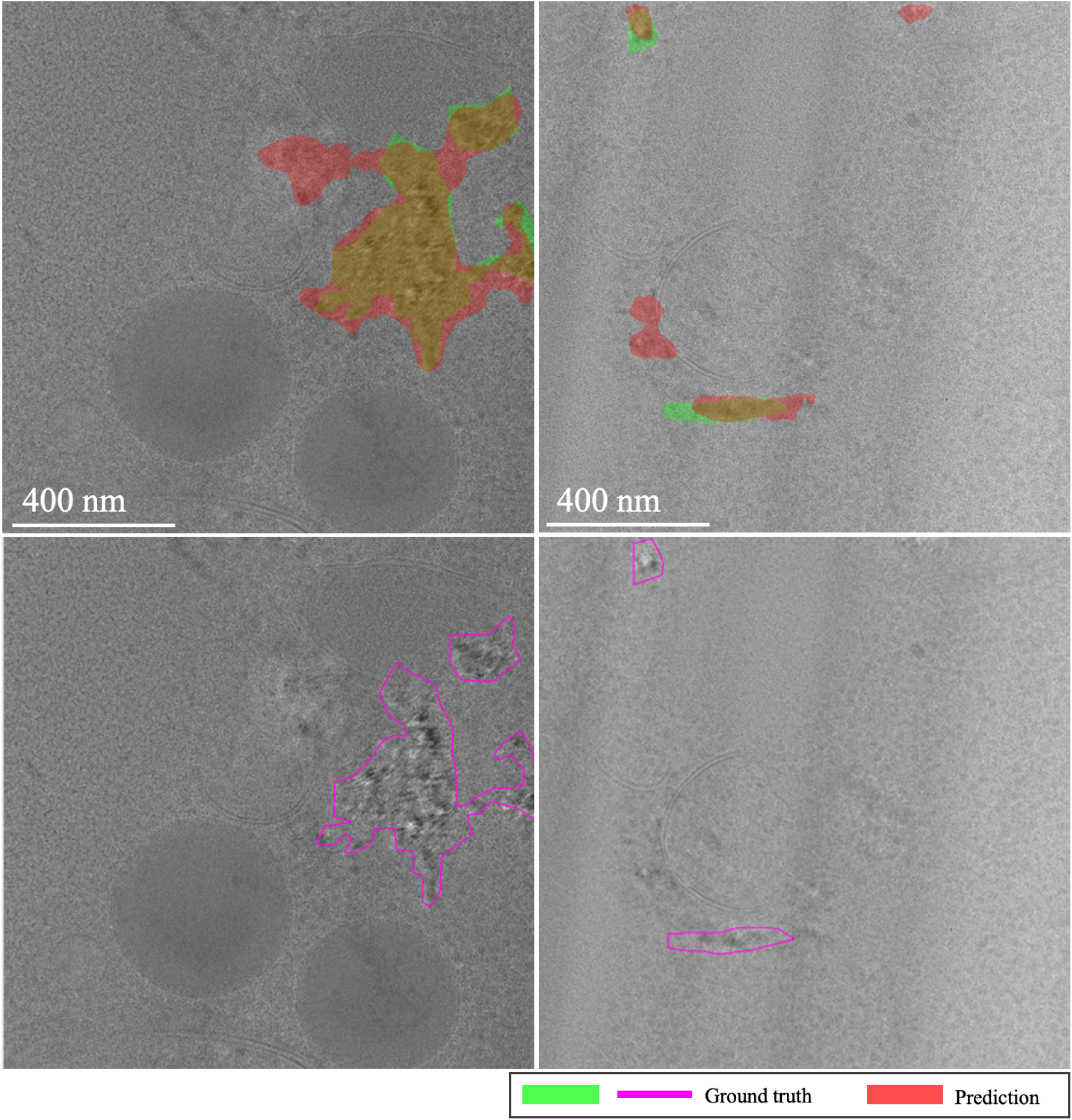
Variability in ice detection in EMPIAR-11166. The images highlight the challenges in detecting less prominent ice features. The first column shows sheet-like patterns, while the second column depicts isolated spots and small regions of low-frequency crystalline ice reflections, demonstrating the models’ adaptability and limitations.

### EMPIAR-11221

Performance was notably lower across models (see Fig. 15). The baseline model showed gradual improvements with some instability, while MAML exhibited volatile behavior due to the diversity and difficulty of the examples encountered during fine-tuning. A primary error source was the significant variety in vitrification combined with a substantial domain shift. Extreme cases illustrated in Fig. 16 show challenges like unvitrified matrices with “brushstroke” reflections, where predictions inaccurately formed extensive ice sheets, leading to substantial mismatches with ground truth annotations (see Fig. 16, first column). Conversely, high-quality samples with minor “brushstroke-like” features were overly conservative, predicting virtually no vitrification issues (see Fig. 16, second column). Additionally, unusual artifacts caused false positives, particularly when contamination was clustered and off-axis, as opposed to when it was unclustered and aligned with the sample, where the model correctly identified and distinguished contamination from ice (see Fig. 17, first and second columns, respectively).

**Figure 15.**
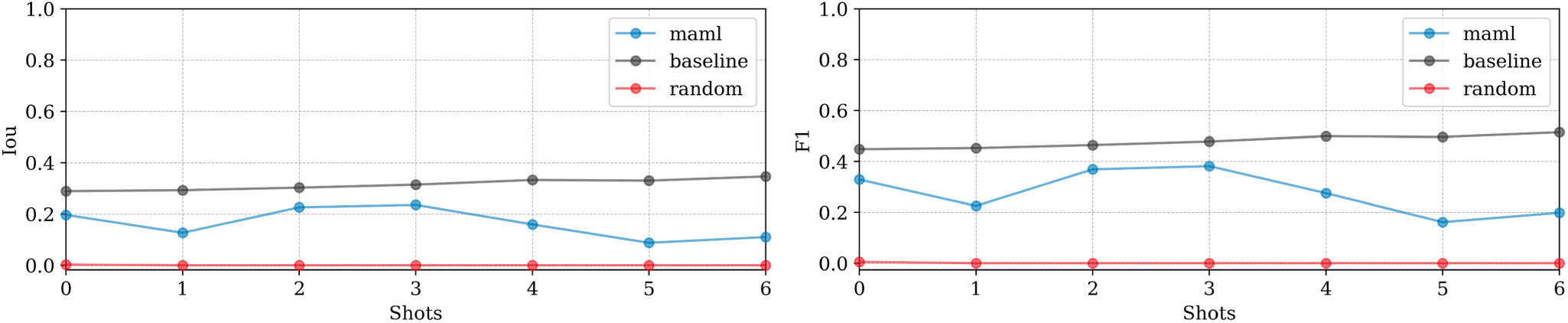
Domain adaptation performance: EMPIAR-11221. The graphs display the performance metrics of different models (MAML, baseline, and random) evaluated using Intersection over Union (IoU) and F1 scores across multiple k-shot scenarios. The performance highlights the challenges of adapting to diverse and difficult examples, with significant domain shifts affecting model accuracy.

**Figure 16.**
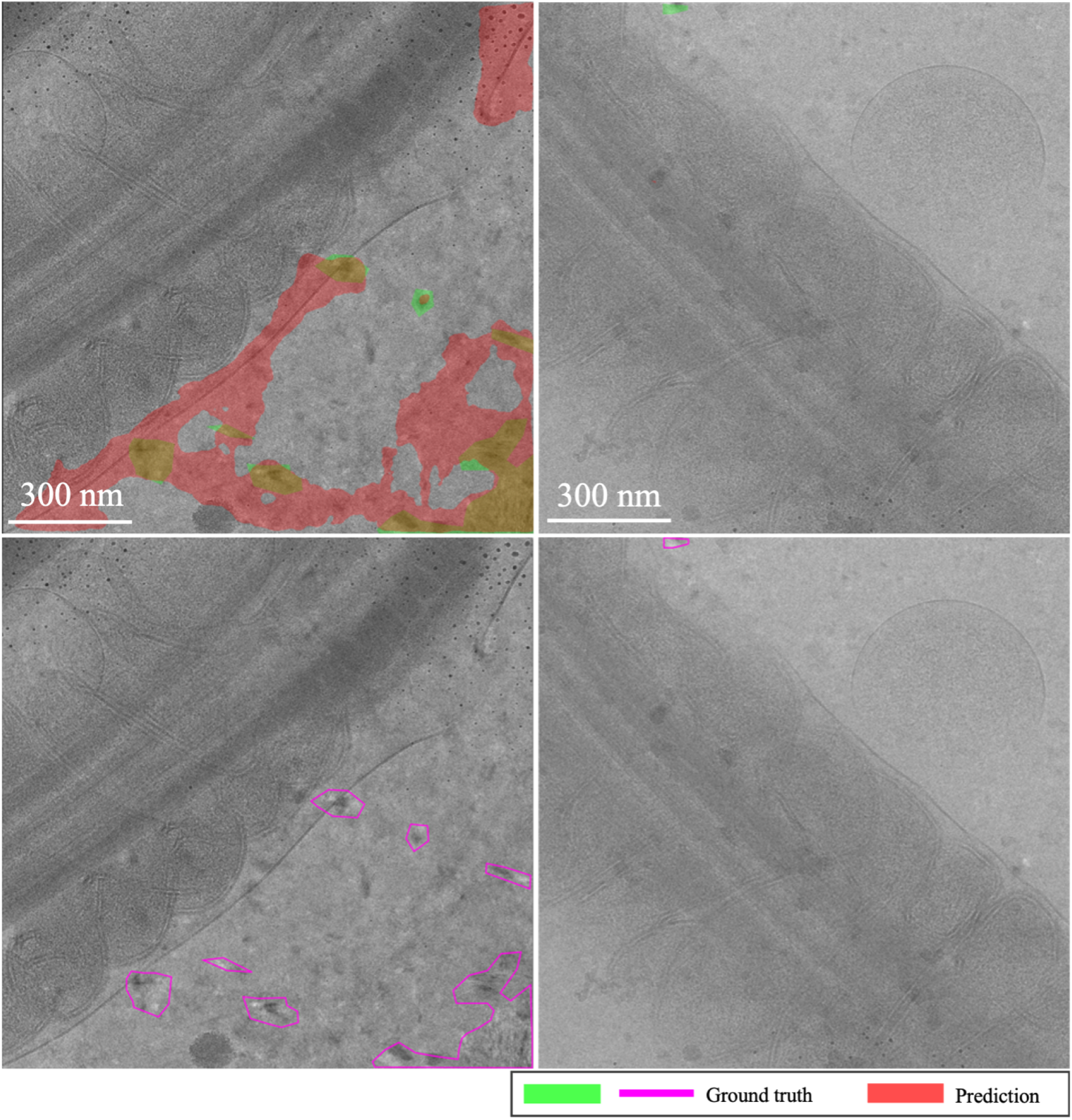
Brushstroke reflections and ice sheet formation in EMPIAR-11221, contrasting extremes from prevalent ice mispredictions to scenarios with no predicted ice issues. The images illustrate the challenges faced by the models in detecting and accurately predicting the extent of vitrification in diverse samples.

**Figure 17.**
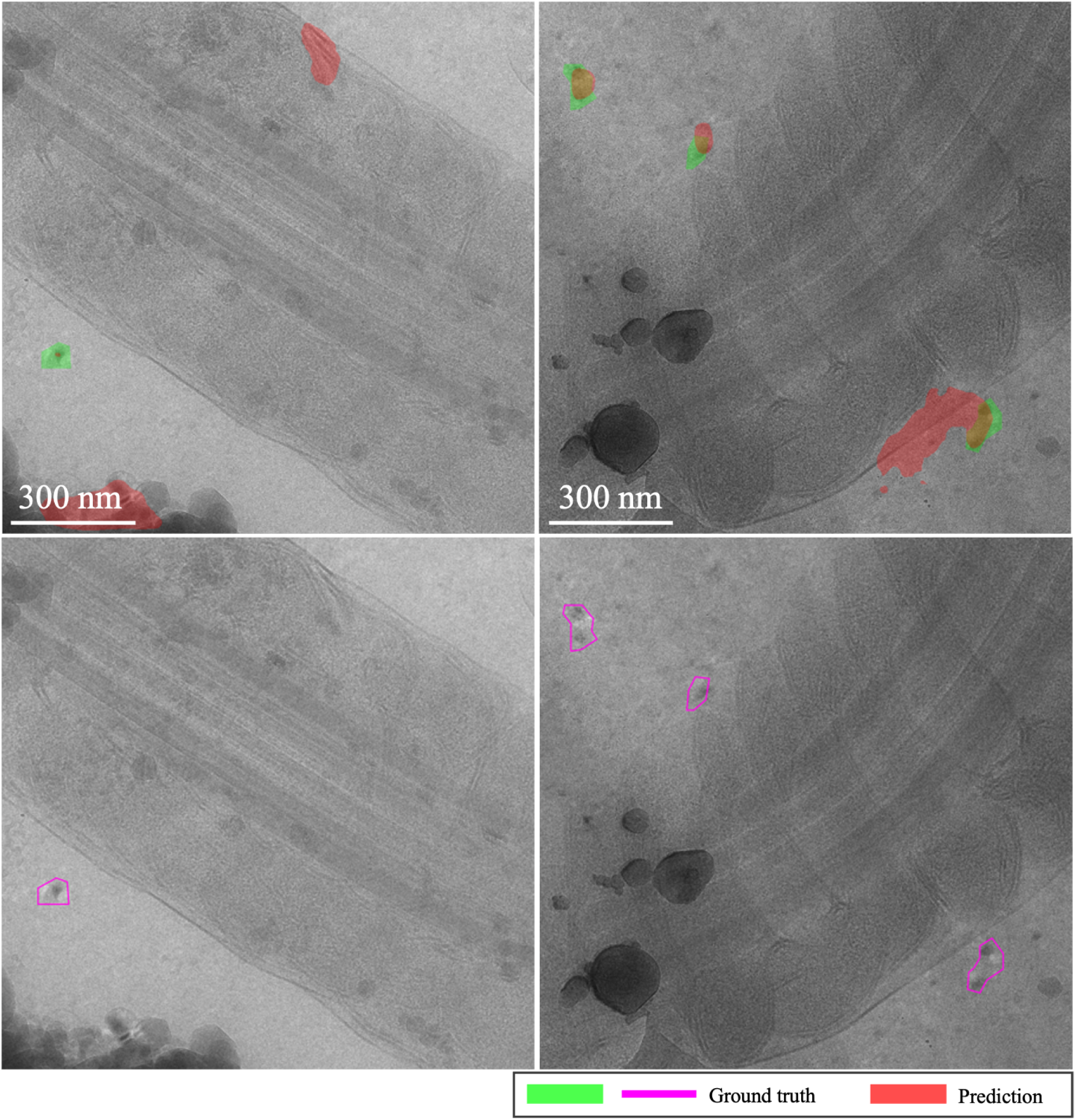
Contamination identification versus ice detection in EMPIAR-11221. The images show the model’s ability to distinguish between clustered contamination and ice, highlighting the difficulties in correctly identifying and labeling these features in challenging conditions.

### EMPIAR-11830

The baseline model exhibited nearly constant performance, indicating a minimal domain shift from the training dataset. In contrast, MAML performance improved smoothly, matching the baseline by the 6th k-shot iteration (see Fig. 18). Challenges included misidentifying sharp chloroplast features as ice patterns, illustrated in Fig. 19. During fine-tuning, examples showcased ice in various phases (see Fig. 20, second column). Nevertheless, at test time it struggled with the recognition of cubic phases alone (see Fig. 20, first column), reflecting a typical class imbalance issue.

**Figure 18.**
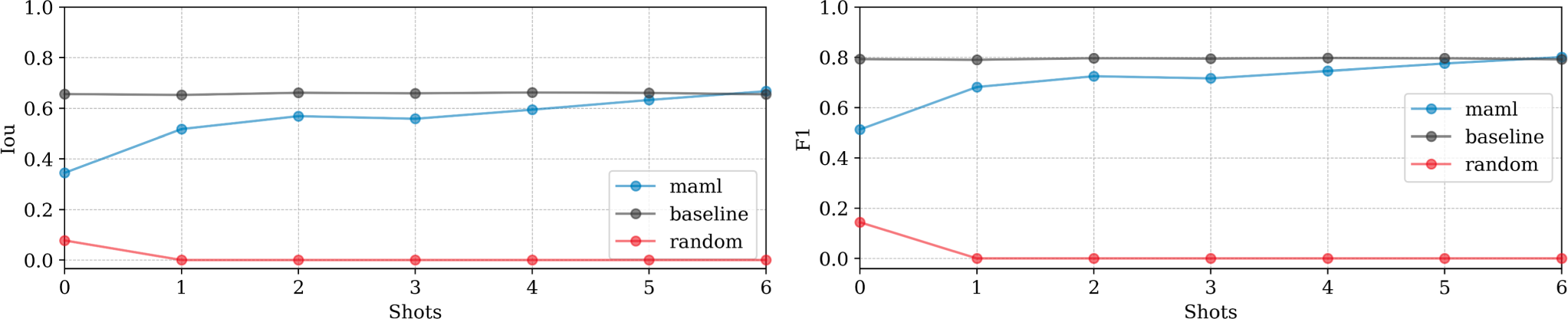
Domain adaptation performance: EMPIAR-11830. The graphs display the performance metrics of different models (MAML, baseline, and random) evaluated using Intersection over Union (IoU) and F1 scores across multiple k-shot scenarios. The MAML model shows a gradual improvement, matching the baseline by the 6th k-shot iteration, while the baseline model’s nearly constant performance indicates minimal domain shift.

**Figure 19.**
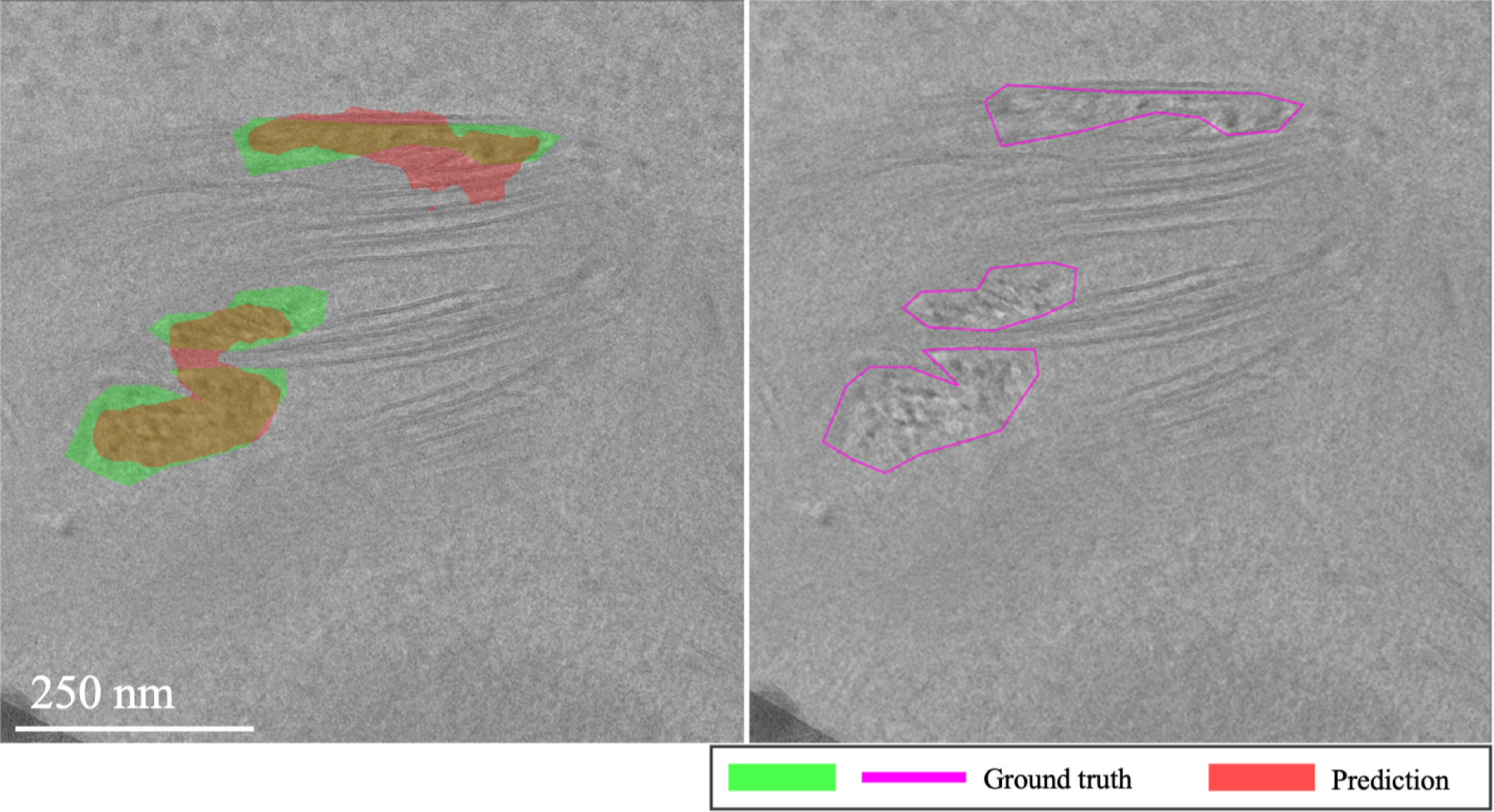
Misidentification of chloroplast features in EMPIAR-11830. The image shows instances where the model incorrectly identified sharp chloroplast features as ice patterns, highlighting the challenges faced in differentiating between these structures.

**Figure 20.**
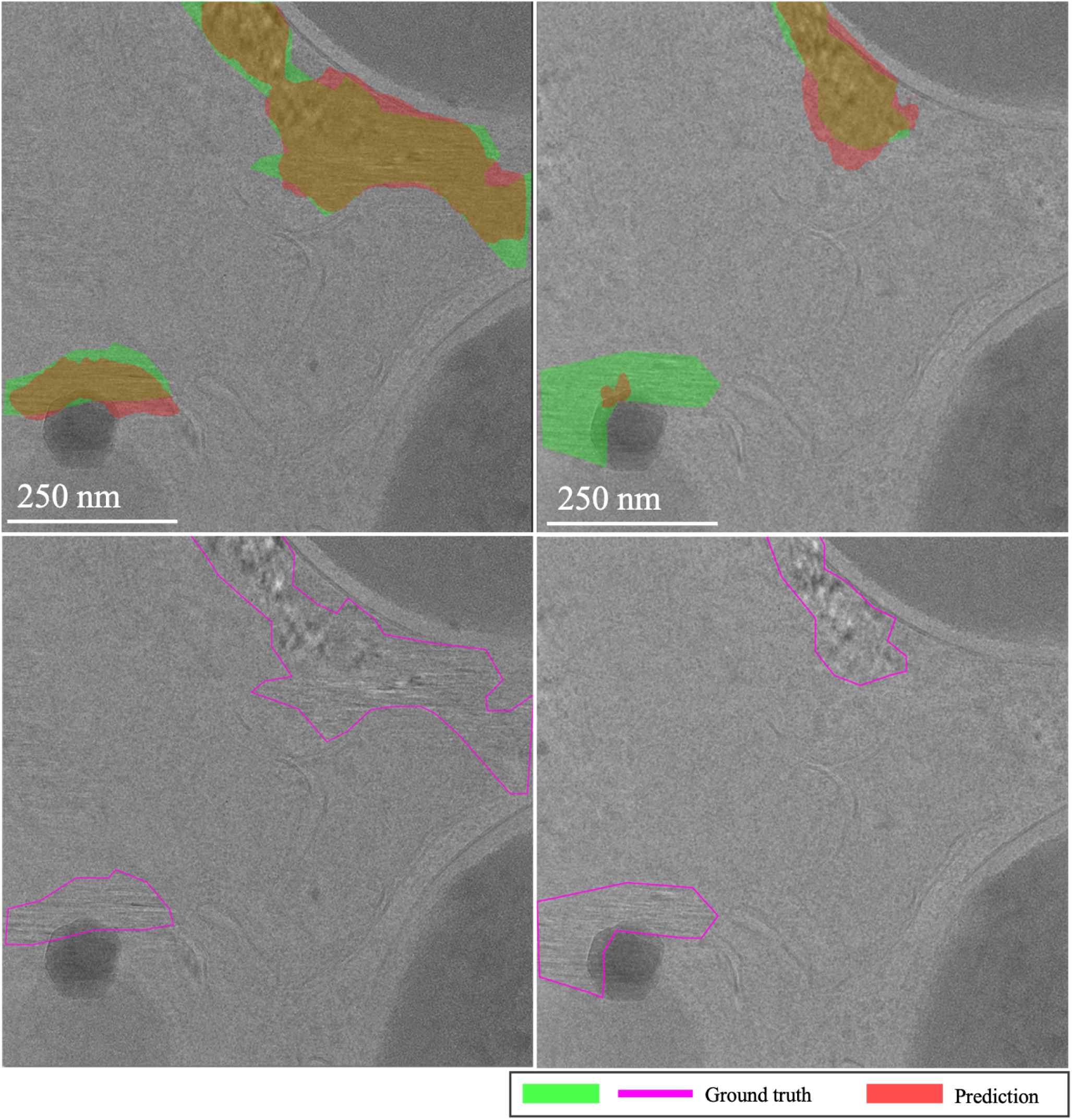
Class imbalance in cubic phase recognition in EMPIAR-11830. The images illustrate the model’s difficulty in recognizing cubic phases during testing, despite fine-tuning examples showcasing ice in various phases. This challenge reflects a typical class imbalance issue where less represented classes are harder to identify accurately.

## Discussion

In this work, we introduced a pioneering approach within the field of cryo-ET by developing a tool that quantifies crystalline ice with precision, filling a significant gap in current methodologies. Our efforts included the first application of the meta-learning paradigm to tomography, effectively redefining multiple tomographic tasks into a unified meta-learning framework. However, constructing a robust and representative dataset emerged as a primary challenge, underscoring the complexity of using a singular distribution dataset for such applications.

Interestingly, while meta-learning presented conceptual promise, transfer learning outperformed it in practical scenarios, proving easier to fine-tune and faster in training. This outcome highlights a crucial point: despite sophisticated algorithmic approaches, inconsistencies in label quality cannot be ignored. To address this, we advocate for the inclusion of temporal dimensions (e.g., LSTM, transformers) in future models to provide additional context, enhancing prediction accuracy and model robustness.

Our Ice Finder tool, optimized for few-shot segmentation, demonstrated the feasibility of adapting quickly to new conditions with minimal examples, offering a user-friendly solution with inference times in the milliseconds range. This capability not only supports the practicality of few-shot learning within this domain but also empowers users to filter tilt series by crystallization percentages, thereby reducing the reliance on manual inspection and significantly streamlining the analytical workflow. Our focus on usability ensures that even simple yet powerful metrics provide clear and actionable insights, enhancing the overall user experience and efficiency.

## Data availability

The tilt series supporting the findings of this study are available in the Electron Microscopy Public Image Archive. The data can be accessed using the following accession codes: EMPIAR-10987, EMPIAR-11058, EMPIAR-11166, EMPIAR-11221, and EMPIAR-11830.

## Code availability

The necessary functions, Jupyter notebook, and best model weights retrained with all data are available at https://github.com/AlmaVivasLago/ice-finder.

## Acknowledgements

We are profoundly grateful to Joshua Soutelo Vieira for his pivotal expertise in meta-learning, which was crucial in the conception and development of this project. Special thanks to Ignacio Arganda-Carreras for his precise feedback on the manuscript, particularly regarding general writing and deep learning concepts. Additionally, we appreciate Oriol Gallego and his team for providing essential data for training, Wanda Kukulski for her expert guidance on sample preparation, and David Gil Carton for his assistance with ice characterization at the microscope. Their collective contributions significantly enhanced the depth and quality of our work.

## Author information

## Author contributions statement

Conceptualization, A.V.L. and D.C.D.; methodology, A.V.L.; software, A.V.L.; validation, A.V.L.; formal analysis, A.V.L.; investigation, A.V.L.; resources, D.C.D.; data curation, A.V.L.; writing—main manuscript text, preparation of figures, review, and editing, A.V.L. and D.C.D.; supervision, D.C.D.

## Additional Information

## Supplementary Information

**Supplementary Table S1:**
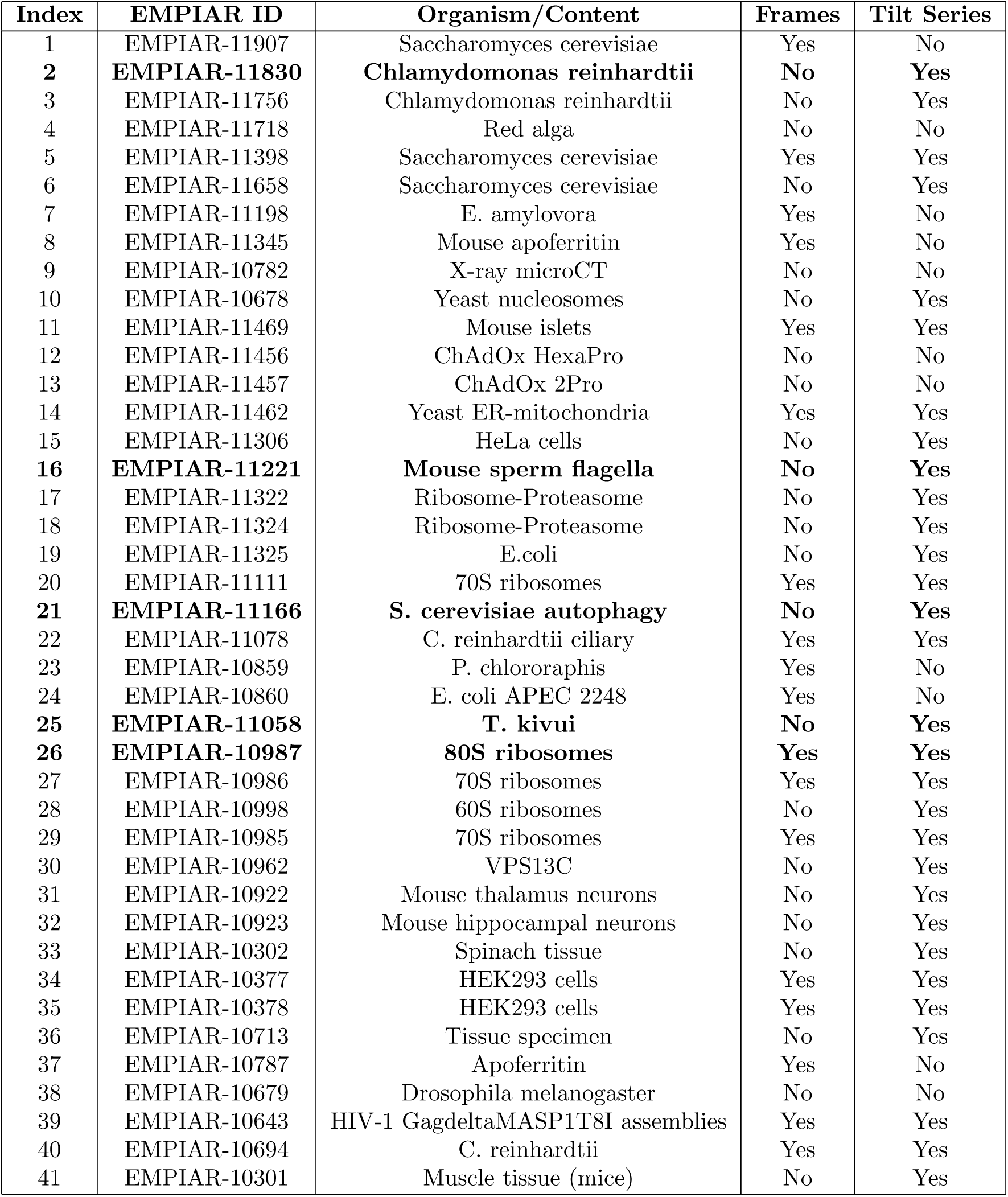
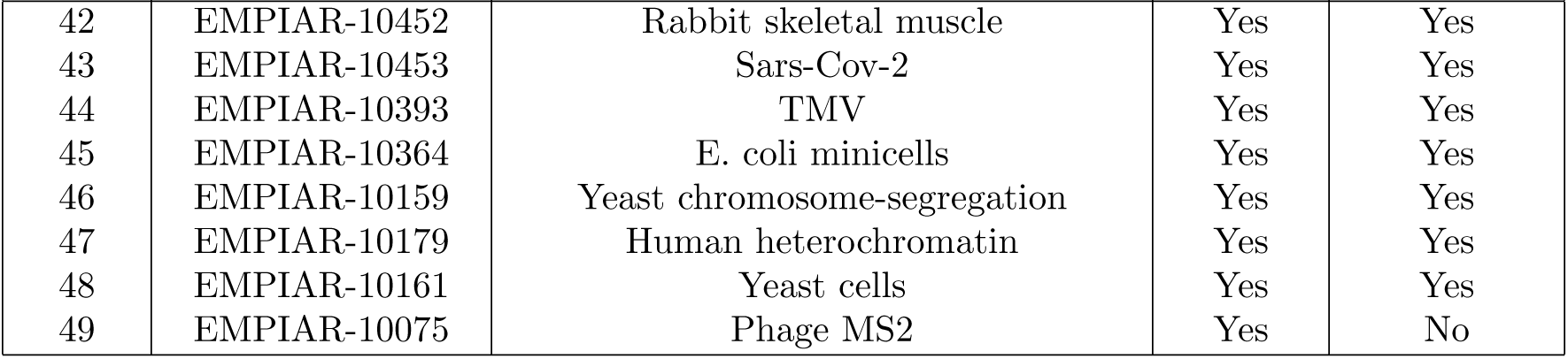
Examples of in situ samples in EMPIAR.

### Implementation and Experimental Details

This section provides a detailed overview of the implementation specifics and experimental configurations utilized in our studies to ensure reproducibility and clarity.

### Computational Setup

Our computational environment included eight NVIDIA GeForce RTX 3090 GPUs and an Intel(R) Xeon(R) Gold 6248R CPU with 96 virtual cores, supported by 500 GB of RAM.

### Software and Frameworks

Experiments were conducted using Python 3.10 and PyTorch 2.0, with CUDA 11.8.

### Training Specifications

Detailed configurations of the models are outlined below.

**Supplementary Table S2:** Training Specifications for Baseline and MAML.

| Parameter | Baseline | MAML |
| --- | --- | --- |
| Steps | 6000 | 25,000 |
| Epochs | N/A | 7 |
| Batch Size | 12 | 2 |
| Learning Rate | 0.001 | Outer: 9.0e-7, Inner: 5.0e-7 |
| Scheduler | CosineAnnealingLR | None |
| Scheduler Parameters $T_{\max}$ | 10 | N/A |
| Optimizer | SGD | Outer: Adam, Inner: SGD |
| Float Precision | 16 | 16 |
| Loss Function | SoftBCEWithLogitsLoss | SoftBCEWithLogitsLoss |
| Input Image Size | 1024x1024 | 1024x1024 |
| **Architecture** | FPN (ResNet34) | FPN (DenseNet121) |
| Encoder Depth | 5 | 5 |
| Decoder Pyramid Channels | 256 | 256 |
| Decoder Segmentation Channels | 128 | 128 |
| Decoder Dropout | 0.2 | 0.2 |
| Input Channels | 1 | 1 |
| Number of Classes | 1 | 1 |
| Activation Function | None | None |
| Upsampling Factor | 4 | 4 |
| Trainable Parameters | 23.1M | 9.3M |
| Estimated Model Size (MB) | 92.596 | 37.174 |

### Reproducibility

Reproducibility was ensured through fixed seeds and controlled environment settings:

~~~
pl.seed_everything(10)
torch.backends.cudnn.deterministic = True
torch.backends.cudnn.benchmark = False
~~~

### Training Durations

Training durations varied significantly across architectures and settings. DenseNet required approximately 11.3 hours under MAML conditions, while ResNet required about 6.629 hours. Baseline models trained completed in 2.45 hours for DenseNet and 1.97 hours for ResNet.

